# Parallel RNA and DNA analysis after Deep-sequencing (PRDD-seq) reveals cell type specific lineage patterns in human brain

**DOI:** 10.1101/2020.04.19.046904

**Authors:** August Yue Huang, Pengpeng Li, Rachel E. Rodin, Sonia N. Kim, Yanmei Dou, Connor J. Kenny, Shyam K. Akula, Rebecca D. Hodge, Trygve E. Bakken, Jeremy A. Miller, Ed S. Lein, Peter J. Park, Eunjung Alice Lee, Christopher A. Walsh

## Abstract

Elucidating the lineage relationships among different cell types is key to understanding human brain development. Here we developed <u>P</u>arallel <u>R</u>NA and <u>D</u>NA analysis after <u>D</u>eep-<u>seq</u>uencing (PRDD-seq), which combines RNA analysis of neuronal cell types with analysis of nested spontaneous DNA somatic mutations as cell lineage markers, identified from joint analysis of single cell and bulk DNA sequencing by single-cell MosaicHunter (scMH). PRDD-seq enables the first-ever simultaneous reconstruction of neuronal cell type, cell lineage, and sequential neuronal formation (“birthdate”) in postmortem human cerebral cortex. Analysis of two human brains showed remarkable quantitative details that relate mutation mosaic frequency to clonal patterns, confirming an early divergence of precursors for excitatory and inhibitory neurons, and an “inside-out” layer formation of excitatory neurons as seen in other species. In addition our analysis allows the first estimate of excitatory neuron-restricted precursors (about 10) that generate the excitatory neurons within a cortical column. Inhibitory neurons showed complex, subtype-specific patterns of neurogenesis, including some patterns of development conserved relative to mouse, but also some aspects of primate cortical interneuron development not seen in mouse. PRDD-seq can be broadly applied to characterize cell identity and lineage from diverse archival samples with single-cell resolution and in potentially any developmental or disease condition.

**Significance Statement:** Stem cells and progenitors undergo a series of cell divisions to generate the neurons of the brain, and understanding this sequence is critical to studying the mechanisms that control cell division and migration in developing brain. Mutations that occur as cells divide are known as the basis of cancer, but have more recently been shown to occur with normal cell divisions, creating a permanent, forensic map of the clonal patterns that define the brain. Here we develop new technology to analyze both DNA mutations and RNA gene expression patterns in single cells from human postmortem brain, allowing us to define clonal patterns among different types of human brain neurons, gaining the first direct insight into how they form.

## Introduction

Although we have learned a great deal about development of the cerebral cortex from animal models, we have remarkably little direct information about how the human brain, which differs vastly in shape, size, and composition from the brains of non-primates, forms the neurons of its cerebral cortex (1–4). Recent studies defining the fundamental cell types of the adult and developing human cortex (5–7) form a foundation for understanding how these cell types develop, how the unique aspects of the human cortex come about, and how developmental brain disorders might alter patterns of cell lineage or cell type in human brain. However, whether individual neural progenitor cells (NPCs) in embryonic stages are restricted to produce certain subtypes of neurons, or multi-potential to generate all neuronal types, is still an open question even in model animal species, since making this distinction requires simultaneous identification of cell lineage and transcriptional analysis of cell type, which remains a technical challenge (8–12).

Somatic genetic mutations accumulate with each cell division during early development, when spontaneous DNA damage escapes the DNA repair machinery, with single-nucleotide variants (SNVs) being the most common mutation type (13–15). The timing of somatic mutations can be inferred by either the cell fraction that carries each mutation or the co-occurrence status of multiple mutations, in which early mutations should be shared by a large fraction of cells whereas later mutations should be present in nested subpopulations of cells (16). Previous study has shown the ability to use somatic SNVs as a rich internal lineage map to birthdate the developmental timing of each neurons differentiated from neuronal progenitor cells (14) but has not combined that with direct analysis of the subtypes of neurons, defined by morphology, location, physiology, or RNA transcription pattern.

Single-cell transcriptomes provide granular information about cell identity (5–7), but it cannot provide lineage maps as it fails to capture most somatic mutations, since somatic mutations occur throughout the genome, most often in intronic or intergenic regions (16, 17). Similarly, DNA-sequencing alone fails to provide information about cell identity, and so lineage mapping using only somatic mutations from DNA sequencing is unable to address questions about the lineage of specific cell identities in neurodevelopment. Somatic mutations in mitochondrial DNA have been recently suggested as potential lineage marks as well, but the modest target size of the mitochondrial genome, and the multiple diverse mitochondrial genomes in each cell, represent challenges to the use of mitochondrial mutations as a rich source of stable lineage markers (18).

To address this challenge, we developed <u>P</u>arallel <u>R</u>NA and <u>D</u>NA analysis after <u>D</u>eepsequencing (PRDD-seq) that identifies somatic SNVs (sSNVs) from single cell and bulk wholegenome sequencing (WGS) data, with multiplexed detection of sSNVs and multiple RNA marker transcripts from single nuclei. We then benchmarked the performance of the DNA and RNA assays of PRDD-seq against bulk WGS and single-cell RNA sequencing (scRNAseq), respectively. Applying PRDD-seq to two postmortem brains of individuals without neurological disease allowed unprecedented quantitative analysis of cell lineage in the human brain. While revealing the expected patterns of divergence of excitatory and inhibitory lineages and “inside-out” generation of excitatory neurons, our PRDD-seq data also directly suggest complex patterns of interneuron formation in the human brain.

## Results

### Simultaneous cell type and lineage analysis of single-cells by PRDD-seq

The workflow of PRDD-seq is illustrated in Figure 1. Single NeuN+ cortical neuronal nuclei from prefrontal cortex (PFC) of postmortem human brain tissue were purified by fluorescence-activated nuclear sorting (FANS) (16) (Fig. 1A), and subjected to one-step RT-qPCR with target-specific primers for 1] cDNA specific for up to 30 marker genes of major neuronal cell types, and 2] specific genomic DNA (gDNA) loci representing identified somatic mutations (see below) as markers of cell lineage (Fig. 1B). Aliquots of the pre-amplified gDNA and cDNA libraries were analyzed for the presence of specific somatic mutations and transcripts by microfluidic genotyping and gene expression profiling, respectively, using the Fluidigm Biomark system (Fig. 1C). The somatic mutations used in PRDD-seq were identified by singlecell MosaicHunter (scMH), described below, a new bioinformatic tool to identify lineage-informative sSNVs, jointly considering WGS data from MDA-amplified single cells and matched deep (>200X) WGS from bulk DNA samples collected from the same brain region (Fig. 1D).

**Figure 1.**
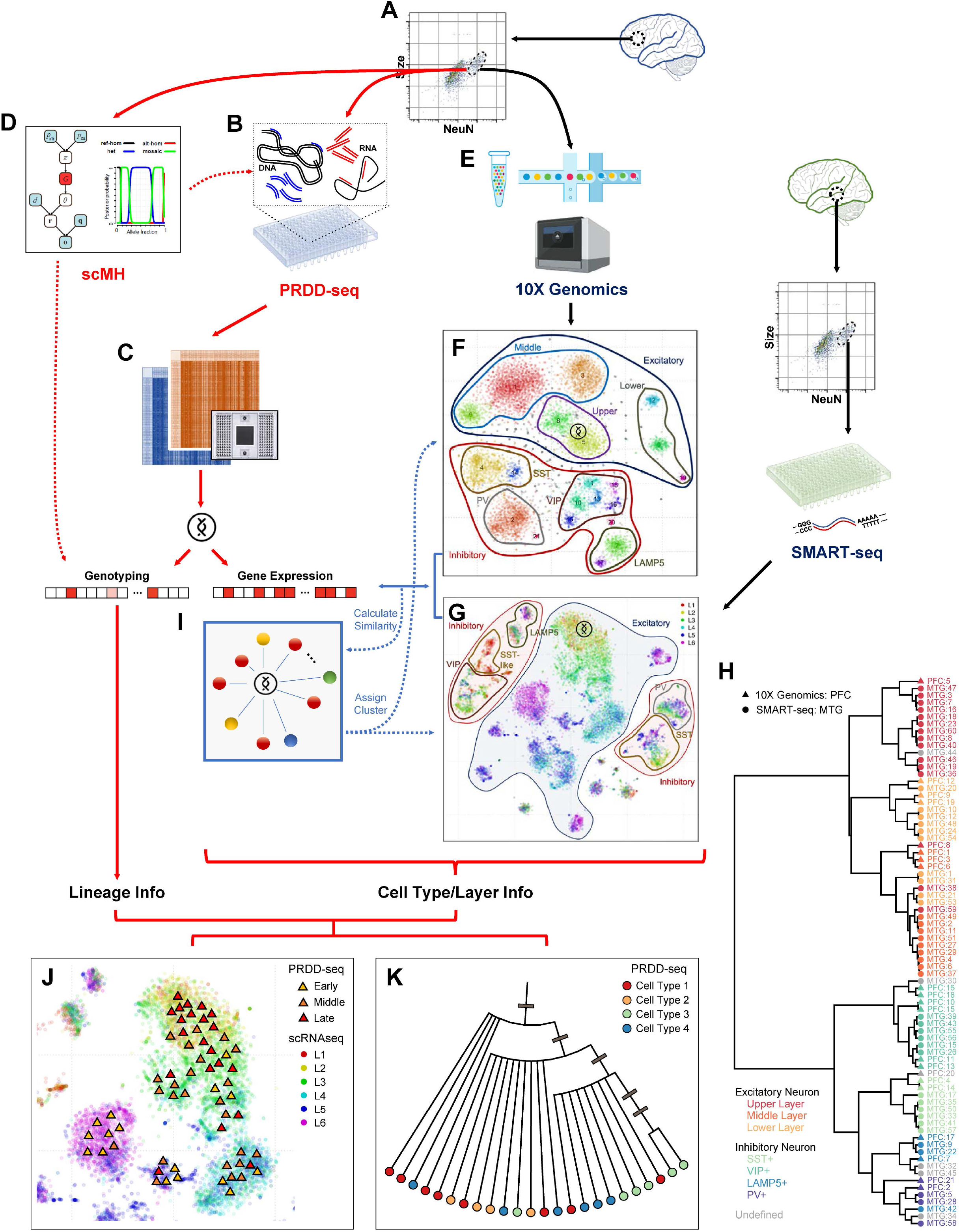
PRDD-seq enables simultaneous assessment of cell identity and lineage in single cells. A. Neuronal nuclei from postmortem human brain were based on NeuN+ immunoreactivity. B. Target-specific one-step RT-qPCR amplification of cDNA and gDNA fragments of interest. C. Single-cell MosaicHunter co-analysis of single-cell and bulk deep sequencing data to identify lineage-informative somatic SNVs. D. Multiplex analysis of the amplified cDNA and gDNA fragments to genotype the somatic SNVs and profile 30 cell type-specific markers of gene expression. E. 10X Genomics scRNAseq was performed on NeuN+ nuclei isolated from the same PFC region. F. 21 cell clusters were identified based on 10X Genomics gene expression data, and then divided into upper, middle, and lower layer of excitatory neurons and four subtypes of inhibitory neurons. G. A second scRNAseq dataset (5) performed on nuclei isolated from the MTG region of another post-mortem healthy human brain was also analyzed where layer information was identified based on layer micro-dissection. Cell types were identified based on gene expression data. H. Transcriptional clustering revealed similar single-cell expression profiles between 10X Genomics PFC and SMART-seq MTG scRNAseq datasets. Cell clusters were color-coded to denote different cell type annotation, and clusters derived from 10X Genomics PFC (triangle) and SMART-seq MTG (circle) in general clustered by cell type but not by platform. I. Each PRDD-seq cell was mapped to the t-SNE maps by the cosine similarity of gene expression to scRNAseq cells, and then assigned cell type and dissected layer accordingly by majority voting of 25 nearest neighbors. J-K. A combination of genotype and gene expression information of PRDD-seq cells allowed lineage and birthdate analysis of particular cell types/layers (J), and cell type differentiation analysis of particular lineage reconstructed by somatic mutations (K). Colored triangles in (I) indicate PRDD-seq cells. Gray bars in (K) indicate occurrences of somatic mutations, whereas all cells in one corresponding sub-clade share the same somatic mutation.

We first created a map of neuronal cell types by analyzing >25,000 single neuronal nuclei -- FANS-sorted based on NeuN immunoreactivity -- by scRNAseq from two different datasets, to create a cell type landscape onto which PRDD-seq analyzed neurons could be located. We performed 10X Genomics scRNAseq of 10,967 NeuN+ nuclei from the same PFC region of one of the brains from which DNA mutations were identified (Fig. 1E). t-SNE analysis of this dataset defined 21 transcriptionally distinct cell clusters, including 8 excitatory neuron clusters that further clustered into upper, middle, and lower layers, and 13 inhibitory neuron clusters that could be further classified into SST+, PV+, VIP+, and LAMP5+ subtypes (Fig. 1F and *SI Appendix,* Fig. S1) (5, 7). A recently published scRNAseq dataset of 15,928 single neuronal nuclei from human middle temporal gyrus (MTG) (5), sorted by NeuN immunoreactivity following microdissection of cerebral cortical layers, provided additional direct information about layer location of neuronal types (Fig. 1G and *SI Appendix,* Fig. S2) and so was used for cell type mapping in parallel. PFC and MTG share relatively generic cerebral cortical architecture as “association” cortex, and clustering analysis of the two datasets (Fig. 1H) shows that they identified similar major cell types, with cells clustering by cell type rather than by platform, although the SMART-seq dataset from MTG defined finer subdivisions of cell type as expected because of its larger sample size and deeper sequence depth.

We jointly analyzed single PRDD-seq cells and scRNAseq cells and mapped each PRDD-seq cell onto the t-SNE maps of scRNAseq based on gene expression similarity (Fig. 1I, see Methods). The cell type and cortical layer information of each PRDD-seq cell was then imputed based on its assigned cluster in scRNAseq datasets. Finally, the combination of genotype and gene expression information of PRDD-seq cells allowed lineage and birthdate analysis of particular cell types (Fig. 1J), as well as analysis of cell type differentiation of particular lineages (Fig. 1K).

### Discovery of lineage-informative sSNVs from bulk brain and single-neuron WGS data

The resolution of lineage reconstruction is dependent on having a comprehensive list of somatic mutations identified from the specific brain under analysis. Whereas deep WGS (e.g., 200-250X coverage) of “bulk” DNA, isolated from tissue, efficiently identifies sSNVs present in 4% or more cells (19), it is insensitive to detecting later-occurring sSNVs that mark late cell lineage events. On the other hand, WGS of DNA amplified from single neuronal nuclei (16) identifies later-occurring sSNVs but is limited by cost and subject to artifacts during single-cell amplification. Therefore, we developed scMH, which incorporates a Bayesian graphic model (20, 21) that integrates analysis of bulk WGS and single-cell WGS data to distinguish somatic mutations from germline mutations and technical artifacts (Fig. 2A; see Methods). scMH first calculates the likelihood and mosaic fraction of candidate sSNVs from a bulk DNA sample, and then applies these values as the priors to genotype each candidate SNV across every single cell being analyzed. The shared presence of a given sSNV in bulk DNA and one or more single cells serves as validation of the sSNV. To expand the utility of scMH when a matched bulk sample is unavailable, we further designed a “bulk-free” mode that can utilize a “synthetic” bulk WGS dataset, generated by *in silico* merging of the many WGS datasets of multiple single-cells obtained from the same donor. We benchmarked scMH using 45X single-cell WGS of 24 neurons—22 of which were sequenced in previous studies (16, 17) —as well as ~200X bulk WGS of PFC (both from the brain of the same individual, UMB1465, who died at age 17 with no neurological diagnosis), against existing single-cell sSNV callers including Monovar (22), SCcaller (23), LiRA (24), and Conbase (25). Sensitivity and false discovery rate (FDR) were estimated based on experimentally validated mutations and clade annotations identified previously (16). With either PFC bulk or synthetic bulk, scMH outperformed the other tools and achieved ~70% sensitivity to detect lineage-informative mutations with < 5% FDR; combining both the default and “bulk-free” modes improved detection sensitivity to 93% without increasing the FDR, suggesting that the “bulk-free” mode of scMH can detect sSNVs that are present in multiple single-cells but may be undetectable in the bulk 200X WGS samples because of the low mosaic fraction of these late mutations (Fig. 2B).

**Figure 2.**
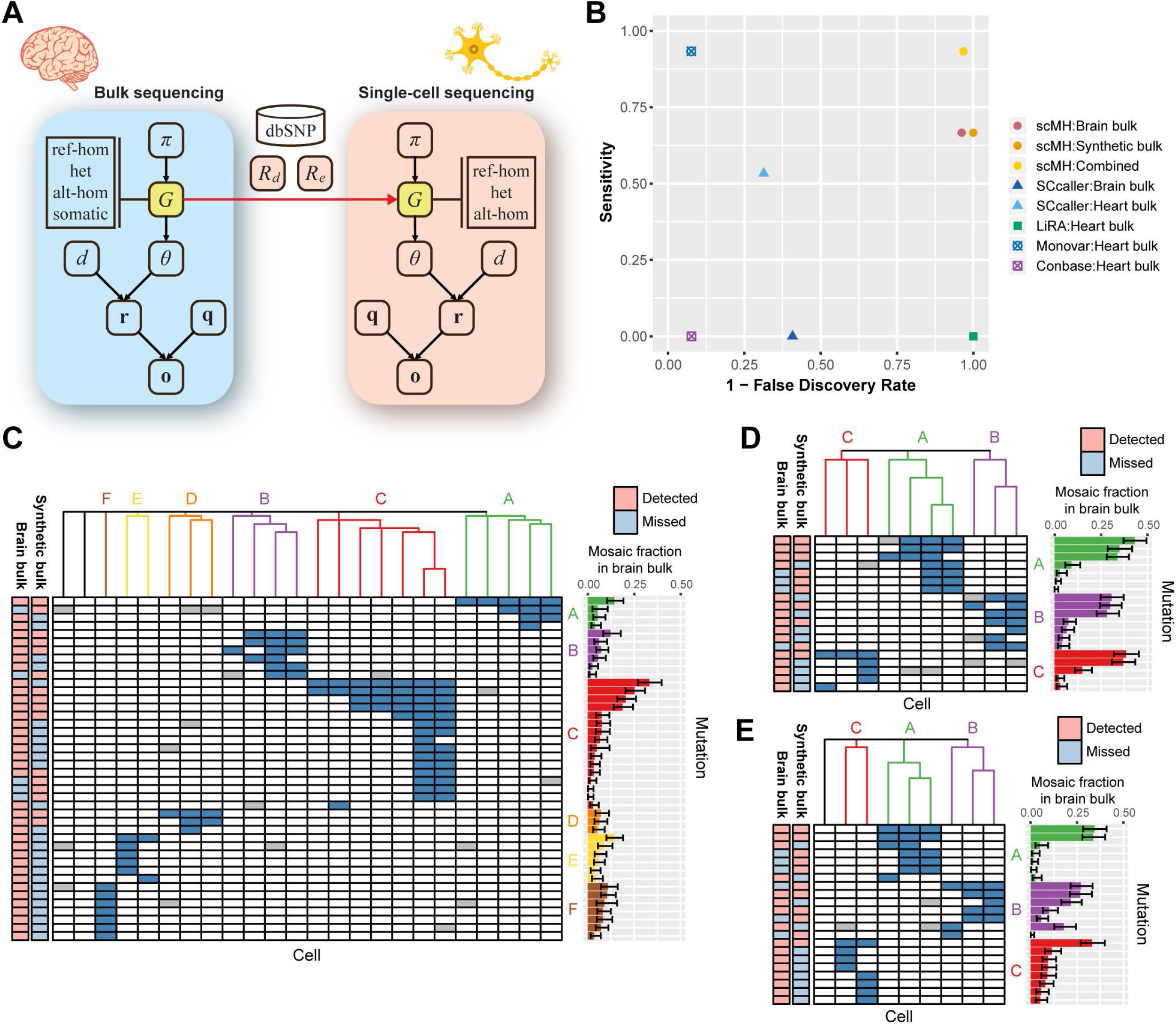
scMH identifies lineage-informative sSNVs from the joint analysis of bulk brain and single-neurons. A. Overview of the extended Bayesian model of scMH to use bulk sequencing data to facilitate sSNV calling in single cells. *G* denotes the genotype state, *π* denotes the prior probabilities of genotype, and *d*, **o, q** denote the depth, observed bases, and their base qualities in bulk or single-cell sequencing data. B. Specificity and precision of identifying sSNVs using scMH and other published callers. scMH outcompeted other callers in both precision and sensitivity. C-E. Validated lineage-informative sSNVs identified by scMH in UMB1465 (C), UMB4638 (D), and UMB4643 (E). Heatmaps demonstrate the genotyping status of sSNVs; dark blue and white squares denote the presence or absence of sSNVs in a given cell, whereas grey squares denote unknown genotype due to locus dropout in single-cell WGS. Bar graphs show the mosaic fraction of each sSNV in WGS of bulk brain sample. Clade E in (C), and clade C in (E), represent likely branching clades where early shared mutations are present, while later sSNVs mark two branches with distinct mutations. Error bars reflect 95% confidence internals.

Applying scMH to data from brains of three normal individuals (UMB1465, UMB4638, and UMB4643 (16, 17), identified and validated 42, 19, and 22 sSNVs, respectively (Fig. 2C-E, and *SI Appendix,* Table S1), with an overall validation rate of 74.8% determined by Sanger sequencing of independently sorted neurons from the same brain region. The number and validation rate of lineage-informative sSNVs detected by scMH dramatically increased from previous studies (16, 17). sSNVs identified from all three brains showed an enrichment in C>T mutations, especially in CpG sites (*SI Appendix,* Fig. S3), a pattern observed in other studies of embryonic mutations and cancer mutations (13, 26), since such C>T mutations appear to be caused by cytosine deamination that is replicated into a fixed SNV before it can be repaired (27). Unsupervised clustering analysis grouped the 24 sequenced neurons from UMB1465 into six different clades; no cells harbored mutations of multiple clades, suggesting the high accuracy of scMH for single-cell genotyping of sSNVs (Fig. 2C). In clades C and E, we observed neurons that shared early mutations but harbored different sets of later mutations, suggesting that they were derived from different branches of the same clades (Fig. 2C). Clustering of ten and nine sequenced neurons from UMB4638 and UMB4643—respectively by their sSNVs— demonstrated similar nested patterns forming three primary clades for each individual and also showed evidence for branches of these clades (Fig. 2D, E). The mosaic fraction of each sSNV in “bulk” DNA (Fig. 2C, D, E) was used as an additional indicator of the sequence in which sSNV occurred, since early sSNVs tend to be found in many single cells, as well as at higher mosaic fraction in bulk DNA, whereas later mutations appear in fewer cells and lower mosaic fraction in bulk DNA. These two findings correlated very strongly.

### Lineage and cell type identity of single-neurons revealed by PRDD-seq

To assess the performance of PRDD-seq in capturing lineage and cell type information from single-cells, we applied PRDD-seq to 1,710 cortical neurons from UMB1465 PFC, using probes to detect 30 out of 42 validated sSNVs in UMB1465, for which we successfully designed highly specific and sensitive probes (*SI Appendix,* Table S1), along with 30 marker genes whose expression levels distinguish major inhibitory and excitatory neuronal subtypes and cortical layers identified in the scRNAseq datasets (5, 7) (*SI Appendix,* Table S2). Overall, PRDD-seq mapped 1,112/1,710 (65%) cortical neurons from UMB1465 PFC into 20 lineage branches and 6 major clades (Fig. 3A). For each major clade, birthdate-ordered lineage branches were inferred from the nested sSNVs, where earlier derived neurons contained fewer clonal mutations, and neurons generated later harbored additional mutations from subsequent cell divisions (16). The nested nature of sSNVs in clades allow cells to be placed into clades using multiple sSNVs, so that cells whose genomes were subject to allelic dropout—which is not uncommon when single cell DNA molecules are amplified—could still be placed into clades based on other sSNV from the same clade (Fig. 3A and *SI Appendix,* Table S1). On the other hand, only 71/1710 (4.2%) neurons contained sSNVs from multiple clades, suggesting a low rate of false positive amplification or sorting of multiple nuclei into single wells in the DNA assay of PRDD-seq (Fig. 3B, upper panel). 527/1710 (30.8%) neurons showed the absence of any sSNVs from the 6 clades; these neurons may be from other clades in which we did not discover sSNV markers (Fig. 3B, upper panel). In PRDD-seq cells, mosaic fractions of sSNVs correlated linearly with the fractions calculated from ~200X bulk WGS, indicating generally unbiased sSNV detection (Fig. 3B, lower panel and Fig. 3C), and allowing confident inference of the developmental sequence of sSNVs according to the nested pattern.

**Figure 3.**
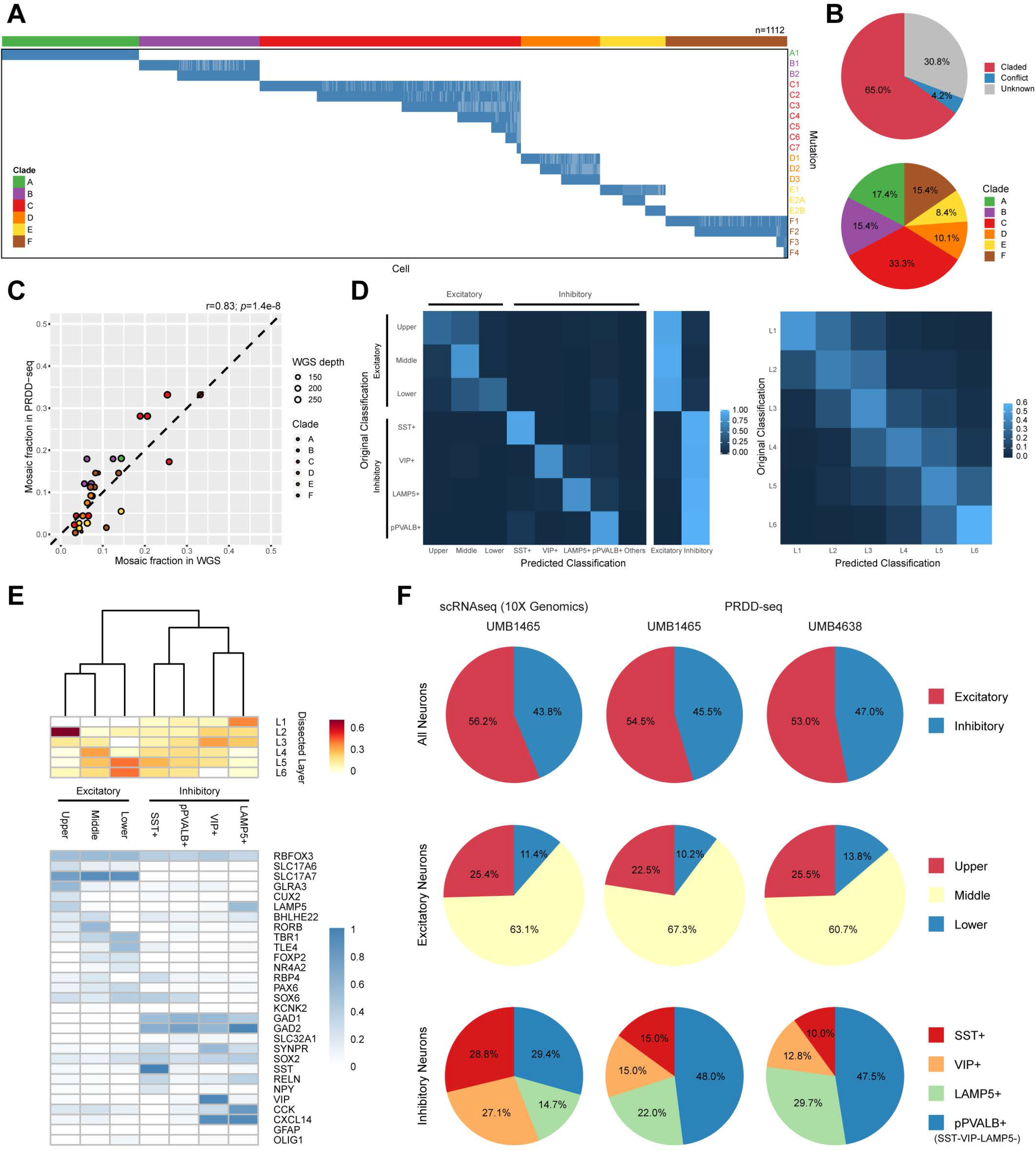
PRDD-seq profiles single-neurons with varied lineage markers and distinct cell type identity. A. Genotyping results of 30 sSNVs (by rows) from 20 lineages across PRDD-seq cells (by columns) from UMB1465. Blue and white squares represent the presence or absence of sSNV respectively, whereas light blue squares represent the sSNVs that were dropouts in PRDD-seq assay but inferred by the presence of deeper mutations from the same clade. B. Clade classification of PRDD-seq cells profiled in UMB1465. In upper panel, PRDD-seq cells which contained sSNVs from multiple or no clades are labeled as “conflict” and “unknown” respectively. C. Correlation of mosaic fractions from WGS and PRDD-seq (calculated as % of assayed cells carrying a given sSNV) in UMB1465. Both methods showed significantly concordant mosaic fractions (Pearson correlation’s *P* < 0.001). D. Accuracy of cell type (left panel) and cortical layer (right panel) classification based on the expression profile of 30 marker genes used in PRDD-seq. scRNAseq cells from each cell type (10X Genomics) and cortical layer (SMART-seq) were randomly sampled and then re-assigned to clusters of t-SNE map using 30 marker genes under PRDD-seq mapping strategy. E. Taxonomy of 3 excitatory layers and 4 inhibitory subtypes based on average expression of 30 marker genes in PRDD-seq cells. Relative density of cortical layers for each subgroup is also shown. pPVALB+ denotes PVALB+/SST-VIP-LAMP5-subtype of inhibitory neurons. F. Relative ratio across different cell types of excitatory and inhibitory neurons between PRDD-seq and 10X Genomics scRNAseq.

Among the 1,112 PRDD-seq cells that were successfully claded, we ran the RNA assay of PRDD-seq to measure the expression of 30 marker genes for each cell. Our evaluation using simulation data derived from our own and published scRNAseq datasets (see Methods) suggested that these 30 marker genes were sufficiently informative to infer many aspects of cell type and dissected layer annotation (Fig. 3D), with an average accuracy of 84% for cortical layer classification (within +/− one-layer difference) and 83% for inhibitory neuron subtype classification. We then utilized expression of these 30 makers to successfully classify 747/1,112 PRDD-seq neurons (67.2%) from UMB1465 into 3 excitatory subgroups—corresponding to upper, middle, or lower cortical layers—and 4 inhibitory subgroups: somatostatin positive (SST+), vasoactive intestinal peptide-positive (VIP+), lysosomal associated membrane protein 5-positive (LAMP5+), and putative parvalbumin-positive (putative PVALB+, or pPVALB+), since probes for PVALB were not always directly assayed (Fig. 3E). PRDD-seq cells assigned to upper, middle, and lower layers by the 10X PFC scRNAseq dataset were also enriched in L2-L3, L4-L5, and L6 markers according to the SMART-seq MTG scRNAseq dataset, respectively, indicating the similarity of the cell type compositions between PFC and MTG, the similarity of the results with the two RNAseq methods, as well as the robustness of the mapping algorithm (Fig. 3E, upper panel). Both our 10X scRNAseq dataset and PRDDseq analysis of UMB1465 and UMB4638 showed higher proportions of inhibitory neurons (43-47%) than reported with other methods, however this ratio was very similar between the three experiments, suggesting that the ratio reflects our particular NeuN+ sorting protocol rather than technical aspects of the cell typing methods (Fig. 3F upper panel). We observed remarkably similar layer and subtype distribution between PRDD-seq and scRNAseq cells for excitatory neurons (Chi-square test; Fig. 3F, middle panel). Among inhibitory neurons, pPVALB+ inhibitory neurons showed a higher proportional representation in PRDD-seq than in scRNAseq, suggesting that a few neurons in this category might reflect amplification failure of the other inhibitory probes (SST, VIP, and LAMP5). In summary, our analysis suggests that PRDD-seq captures the major aspects of cell types, without systematic loss of any given cell type.

### Early divergence of progenitors for excitatory and inhibitory neurons

The simultaneous analysis of lineage and gene expression from the same neurons enabled us to study the change of cell type contribution during early neurogenesis. Using PRDD-seq, we profiled >2700 neurons from two brains, UMB1465 and UMB4638, and successfully captured both lineage and cell type information from 747 and 480 neurons, respectively. In both UMB4638 and UMB1465, all lineage clades showed early sSNVs in both excitatory and inhibitory neurons, reflecting mutations occurring during early embryogenesis before the divergence of these cell types, whereas late SNVs show progressive restriction to one or the other cell type (Fig. 4A, B). Among the six major clades in UMB1465, clade C contained seven nested branches with mosaic fractions diminishing from 0.33 to 0.0067 (Figure 3A and *SI Appendix,* Table S1), with an increasing percentage of excitatory neurons containing mutations C1 to C5, and only excitatory neurons containing mutations C6 to C7 (Fig. 4A), while clade F showed similar progressive restriction. Similarly, both clade A and B in UMB4638 showed nested mutations that became progressively limited to excitatory neurons (Fig. 4B). Interestingly, the excitatory neurons appeared exclusively in branches with mosaic fraction below ~0.04 (Fig. 4A, B, and *SI Appendix,* Table S1), corresponding to a progenitor giving rise to about 4% of the total cells in that cortical sample. Considering that ~40% of cortical cells are excitatory neurons, with the remainder being glial cells or inhibitory neurons (28, 29), this observation suggests that ten or more excitatory neuronal progenitor cells (NPCs) generate excitatory neurons in a given cortical area, or “column”; the fact that 6-7 (including a branched clade) excitatory precursors are explicitly marked by non-overlapping clades, and account for 60-70% of excitatory neurons in our sample, independently supports this estimate. On the other hand, two clades (clade A and B) from UMB1465 are statistically enriched for inhibitory neurons (two-sided one-proportion Z-test’s *P* < 0.05), with the percentage of inhibitory neurons increasing from B1 to B2 (Fig. 4A). These results show that at least some human NPCs demonstrate restricted cell type output, supporting the model first established in mice (30–32) and strongly supported by conserved gene expression patterns in the ganglionic eminence between humans and non-humans (33, 34), that excitatory and inhibitory neurons are generated from distinct progenitor regions.

**Figure 4.**
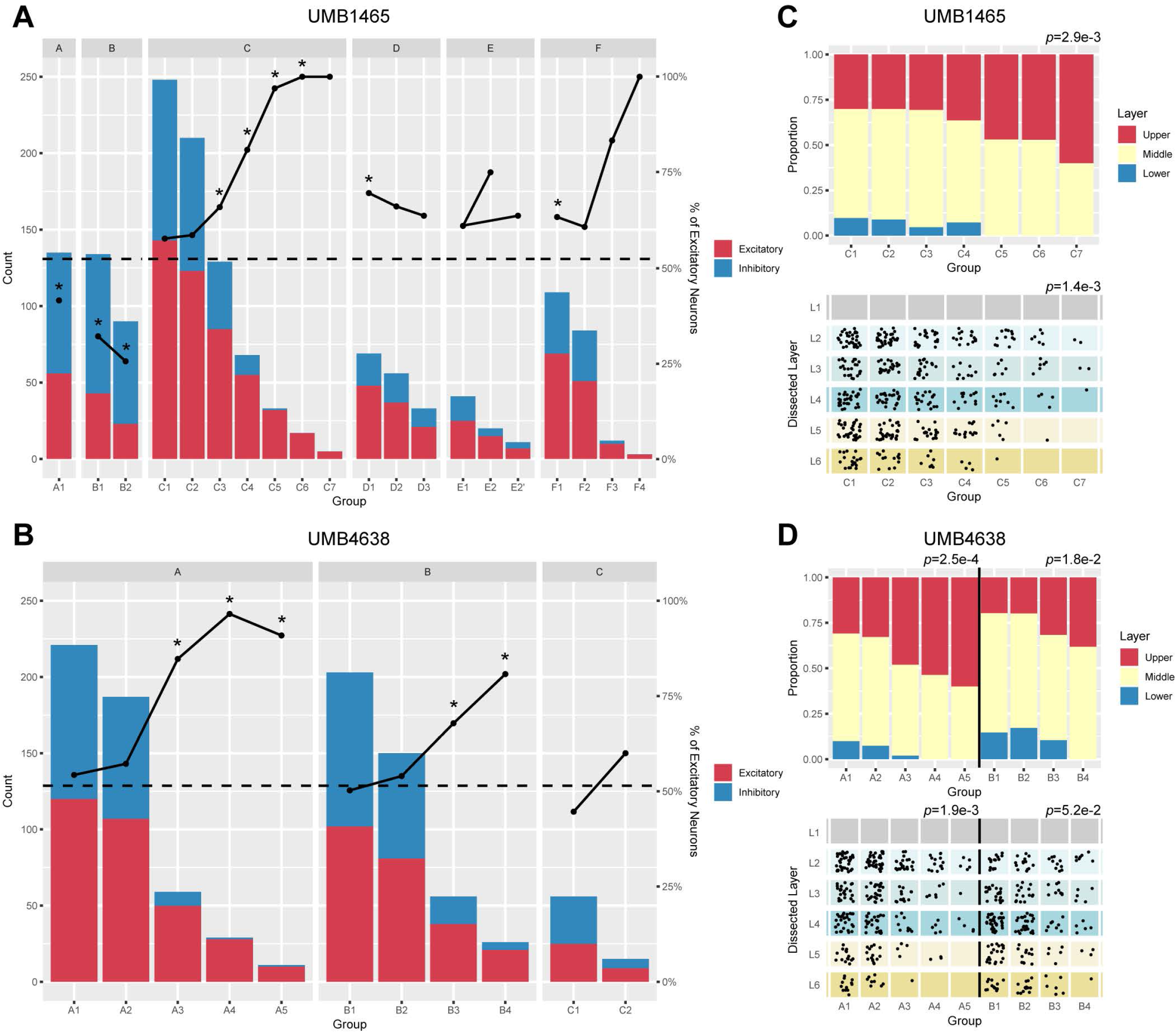
PRDD-seq reveals distinct developmental sequence of excitatory neurons in different cortical layers. A-B. The total number (bar plot) and ratio (dot plot) of excitatory and inhibitory neurons in different lineage clades defined by one or more sSNVs in UMB1465 (A) and UMB4638 (B). Percentage of excitatory neurons increased in later lineage timepoints in clades C and F in UMB1465 and clades A and B in UMB4638. In Clade E of UMB1465, E1 branches into two subclades E2A and E2B. Dashed line: average excitatory neuron percentage. Asterisk denotes significantly different excitatory-inhibitory ratio from the average (two-sided one-proportion Z-test’s *P* < 0.05). In clades C and F from UMB1465, and clades A and B from UMB4638, later mutations become progressively limited to excitatory neurons. C-D. Layer distributions of excitatory neurons in representative excitatory lineages in UMB1465 (C) and UMB4638 (D), respectively. Layers are determined by mapping PRDD-seq cells onto human PFC scRNAseq (upper panels) or human MTG scRNAseq (lower panels) based on the expression profile similarity of marker genes. In all three illustrated clades, the percentage of upper layer neurons increased while that of lower layer neurons decreased in cells containing sSNVs present at lower mosaic fraction. *P*-value was calculated by Pearson correlation with ordinal variables.

### “Inside-out” order of cortical layer formation for excitatory neurons

Further sub-typing of excitatory neurons using laminar markers revealed layer-specific patterns of excitatory neuron neurogenesis. For example, in UMB1465, the percentage of lower layer neurons carrying a mutation decreased from mutations C1 to C4, and no deep-layer neurons were detected carrying C5 to C7, with the percentage of upper layer neurons increasing correspondingly from C1 to C7 (Pearson correlation’s *P* = 2.9×10^-3^; Fig. 4C, upper panel). To gain more precise layer identities of PRDD-seq cells, we mapped them to the SMART-seq MTG scRNAseq dataset obtained after layer microdissection using the same methods as earlier (5), which generated similar “birthdate” patterns in clade C, with early lineage sSNVs present in all layers, and later sSNVs restricted to middle and upper layers (Pearson correlation’s *P* = 1.4×10^-3^; Fig. 4C, lower panel). A similar trend was also observed in clades A and B in UMB4638. Mapping PRDD-seq cells of UMB4638 to both 10X PFC and SMART-seq MTG scRNAseq datasets showed that cells with later lineage markers were restricted to middle and upper layers (Fig. 4D). These results together directly indicate that human cortical excitatory neurons are formed in “inside-out” sequence after preplate cells are born, similar to mouse and non-human primates (35–37). Furthermore, it suggests that neurons in lower cortical layers begin becoming postmitotic relatively quickly after progenitors are specialized for excitatory neuron production.

### Diverse spatiotemporal patterns of development of inhibitory neuron subtypes

Mapping PRDD-seq cells onto two different scRNAseq datasets also allowed analysis of cortical inhibitory neurons, which originate from multiple developmentally transient structures of the ventral telencephalon, including the medial, lateral and caudal ganglionic eminences (MGE, LGE, and CGE), and migrate into dorsal cortex (30, 38). However, the highly dispersed nature of inhibitory neuron clones observed in animal models (39–41) suggests that sSNVs in the inhibitory lineage are likely to be present at exceedingly low allele frequencies in bulk DNA and tiny fractions of single cells, so that only sSNVs occurring relatively early in development have been analyzed so far. Inhibitory neurons derived from MGE and CGE can be distinguished by expression of specific markers (5, 6), and PRDD-seq analysis showed that interneurons with diverse marker genes were generated over the same developmental window (Fig. 5A, B). The analyzed sSNVs were shared by multiple inhibitory subtypes, with hints that late marks might be more limited to cell types, but no differences that reached statistical significance (FDR-adjusted Chi-square test’s *P* > 0.05). Previous studies cataloging interneurons in mouse and human have suggested that MGE-derived inhibitory neuron subtypes (SST+ and PVALB+) are enriched in infragranular cortical layers, while CGE-derived interneuron subtypes (LAMP5/PAX6+, VIP+) tend to occupy upper cortical layers preferentially (5, 42, 43) and thus our mapping of PRDD-seq cells onto scRNAseq reflected these patterns. Birthdating analyses in mice and non-human primates have reached contradictory conclusions about whether inhibitory neurons follow inside-out patterns of generation similar to excitatory neurons (44, 45), though recent analyses in mice suggest that previous contradictions may reflect the convolution of multiple patterns of generation that may be subtype specific (46). We found that MGE-derived pPVALB+ subtype neurons, enriched in layer IV-VI, showed if anything a trend for the latest-generated neurons to show markers of deeper layers (Fig. 5C, D). SST+ neurons, widely distributed in layer II-VI, similarly did not show an inside-out pattern detectable with the mutations and cells analyzed (Fig. 5C, D). We robustly detected SST+ neurons with expression of layer I markers in human PFC (SST-like subclass) (Fig. 5C, D), consistent with observations in MTG (5, 47) and in mice, where such layer I SST+ expressing cells are rare but present (43, 47). These upper layer, CGE-derived SST-like cells are a subclass of LAMP5+ interneurons that are more transcriptionally related to VIP neurons than MGE derived SST+ interneurons, though they lack VIP expression (5, 47). Our data further confirm that LAMP5+ interneurons express markers suggesting broad laminar location, but also did not reveal a simple inside-out progression of formation (5). Interestingly, we observed a substantial proportion of LAMP5+ inhibitory neurons, particularly the SST-like class, labeled by later mutations, indicating that this subtype may be generated later during development than other inhibitory cell types (Fig. 5C, D). Overall, our findings suggest little evidence of the inside-out patterns of neurogenesis demonstrated by excitatory neurons, but also show that detailed analysis of interneurons will likely require deep datasets of sSNV occurring at late stages of interneuron development, and higher-throughput methods of analysis.

**Figure 5.**
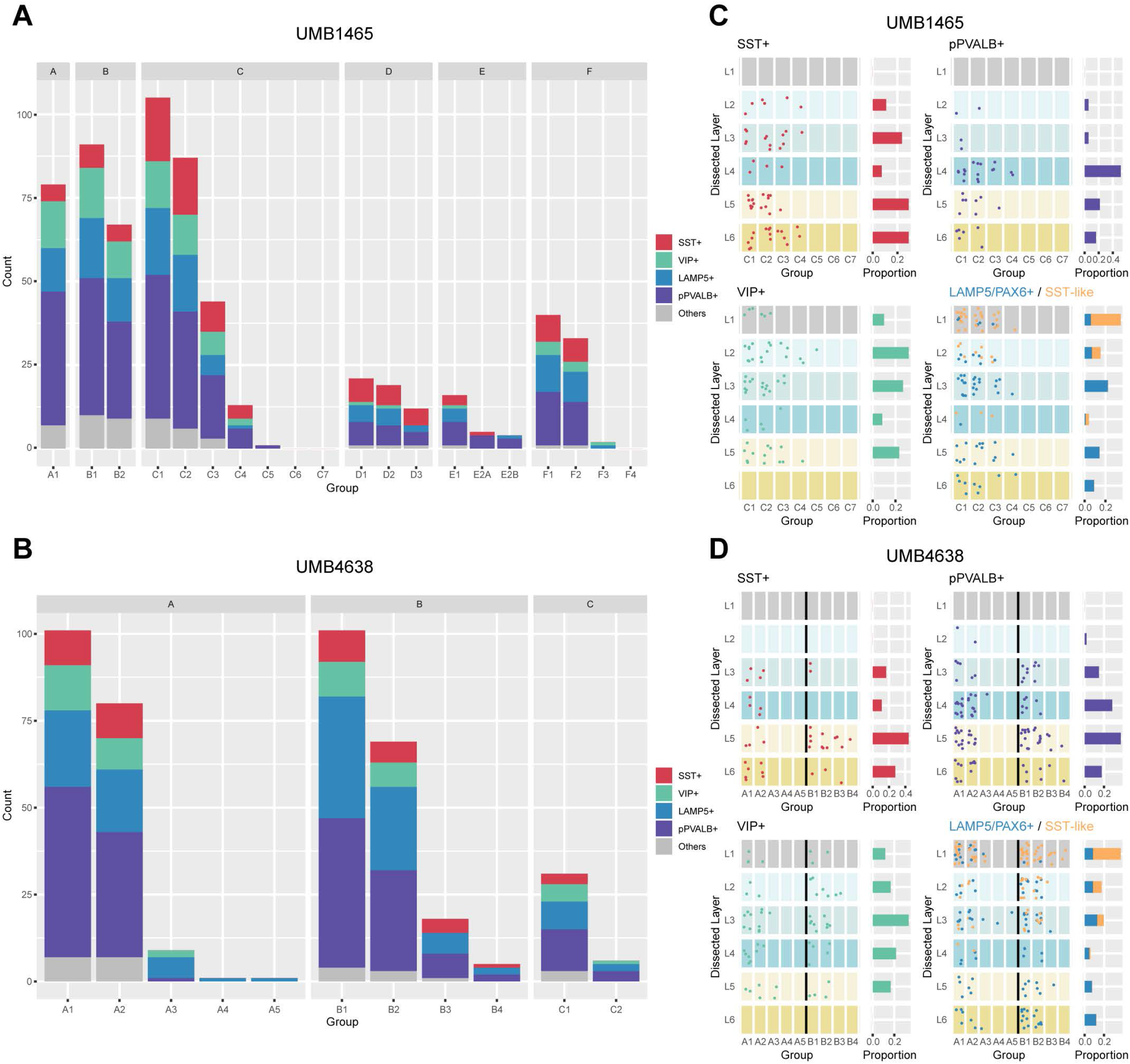
PRDD-seq reveals heterogeneous developmental process for inhibitory neurons. A-B. Distribution of different subtypes of inhibitory neurons in different lineages in UMB1465 (A) and UMB4638 (B), respectively. Major subtypes of inhibitory neurons are widely distributed in different lineages. C-D. Layer distributions of inhibitory subtypes in representative lineages in UMB1465 (C) and UMB4638 (D), respectively. Bar graphs show the proportion of each subtype of neurons in different layers. MGE derived (SST+ and pPVALB+) and CGE derived (VIP+, LAMP5/PAX6+, and SST-like) interneurons showed similar mutation profiles, suggesting that the groups are produced simultaneously. pPVALB+ subtype neurons were enriched in layer IV-VI, while MGE-derived SST+ interneurons showed a similar laminar distribution as pPVALB+ interneurons, with no clear evidence of an “inside-out” birth dating pattern. CGE-derived interneurons were broadly distributed across cortical layers, with SST-like cells heavily favoring supragranular layers; LAMP5+, including SST-like cells, were enriched for later lineage marks, suggesting they may be produced later in development than other subtypes.

## Discussion

We have developed scMH and PRDD-seq that allowed us, to our knowledge, the first simultaneous analysis of cell lineage and transcriptional cell type in human brain—and potentially, any mammalian brain—through improved identification of sSNVs in deep bulk and single-cell sequencing data. Our analysis of a single cortical area (PFC) in two individual brains revealed some conserved patterns of cell lineage compared to nonhumans, including that inhibitory and excitatory neurons diverge early in humans, and that excitatory neurons form following a similar “inside-out” order as seen in the animal models. However, PRDD-seq also provides the first quantitative estimate in any species of number of progenitor cells (approximately 10) that generate the excitatory neurons in a given cortical area. Furthermore, PRDD-seq also provided some direct insight into inhibitory neuron development in humans, supporting parallel development of different subtypes of inhibitory neurons, with spatial and temporal associations specific only to some subtypes. Our data show that, as methods improve to capture sSNVs present in small numbers of cells, the natural occurrence of sSNVs with each cell division (13, 14, 17) is likely sufficient to provide a very rich map of cell lineage patterns in any given postmortem human brain.

The human cerebral cortex has been thought to contain approximately 80% excitatory glutamatergic neurons and 20% GABAergic interneurons (48), although recent scRNAseq studies have reported a somewhat lower ratio of about 70% excitatory neurons (*SI Appendix,* Table S3) (5, 49, 50). Although our PRDD-seq analysis showed 661 excitatory versus 566 inhibitory PRDD-seq cells in total for UMB1465 and UMB4638, which represents 54% excitatory neurons (*SI Appendix,* Table S3), this higher proportion of inhibitory neurons seems to reflect either aspects of the tissue (which was stored for long periods frozen), or our NeuN+-sorting method, since similar ratios are seen in 10X scRNAseq from the one brain analyzed (*SI Appendix,* Table S3). On the other hand, PRDD-seq cells are studied as containing at least one sSNV identified from scMH using a small number of deeply sequenced neuronal nuclei isolated from the same region, and so do not represent an unbiased sampling of the human brain region. Nonetheless, the fact that we can assign 60-70% of all excitatory neurons to clades in UMB1465, and that neurons with identified SNVs represent most major neuronal types in scRNAseq (Fig. 3E), suggests that our sampling has captured the majority of the lineage of the cortical patch, although rare lineages are likely to be missed without much deeper sequencing. Moreover, the presence of 6-7 explicitly marked clades, and the ability to correlate the allele frequency of a sSNV to the excitatory-restriction of the cells carrying that sSNV, allows two independent quantitative assessments of how many progenitors (approximately 10) contribute to the neurons of the patch of cortex from which neurons were isolated, illustrating the remarkable quantitative potential of this approach.

Since occasional dropout of DNA marks and RNA markers in PRDD-seq is unavoidable, limited by the quality of isolated nuclei, we emphasize that our results are most robust when analyzing cells positive for both. The quality of postmortem brain tissues can influence the integrity of both genomic DNA and mRNA. Regarding DNA, since no whole-genome amplification is performed prior to targeted pre-amplification, only a single molecular copy of each allele is available for genotyping of each sSNV, so occasional dropout is inevitable. However, our lineage strategy is based not only on the presence of clade-specific sSNVs but also the absence of many sSNVs from other clades (Fig. 3A), so the chance for mis-assigning cells should be relatively small. Nevertheless, mapping our sSNVs onto our scRNAseq dataset suggests that lineage marks are present in the major neuronal subtypes, although rare neuronal types are likely to be missed given our modest sample size. Regarding RNA, single nuclei from postmortem human brain contains only a small amount of mRNAs. Fluidigm Biomark assays are microfluidics-based qPCR assays that are sensitive to subtle changes of the input or environment. As a result, we observed a 30.4% dropout rate of DNA markers and similar level of dropout of RNA marker dropout. However, since PRDD-seq analyses excluded these dropout events, and were completely based on the relative cell type proportions across different stages within one lineage, we have no reason to think that the dropouts are systematic with respect to cell type with one exception: the relatively larger proportion of pPVALB+ neurons in PRDD-seq than scRNAseq, likely reflecting the failure of some probes for SST, VIP, and LAMP5. Better and richer probe sets are likely to be able to resolve this in the future.

There are limitations to our analysis, since we are analyzing a small sample of the vast size of the human brain, and PRDD-seq is relatively low-throughput and expensive, so our initial analysis only can make conclusions about relatively common cell types. The present analysis is somewhat limited in the analysis of late mutations present in 1% of cells, especially interneurons, since it is challenging to detect those mutations with great sensitivity, but will await single-cell studies on subtypes of neurons in the future. On the other hand, the combined analysis of sSNVs and cell types is archival and progressive. The vast size of the human brain means that each subsequent round of DNA sequencing—whether of bulk tissue or of single or pooled cells—adds to the total depth of sequence data, and provides progressively richer information about late sSNVs. Indeed, the likely dispersed nature of inhibitory clones suggests that analyzing one cortical region could provide sequence data useful in the analysis of a completely different cortical region for these cell types.

Overall, PRDD-seq has many advantages even beyond the quantitative analysis of lineages and mosaic fractions that we begin to illustrate here. Since the method uses sSNVs as lineage marks, it is inherently genomic and so allows correlation not only of normal developmental patterns, but would immediately capture alterations to lineage patterns caused by function-altering germline or somatic mutations. In addition, since sSNVs serve as *in vivo* cellular markers for drawing a developmental lineage map without any transgenic manipulation as demonstrated in this study, the method promises to be applicable in principle to any species or human disease condition for which post-mortem brain is available.

## Materials and Methods

### Human tissues whole-genome sequencing

Frozen post-mortem tissues from three neurologically normal individuals, UMB1465 (a 17-year-old male), UMB4638 (a 15-year-old female), and UMB4643 (a 42-year-old female), were obtained from the NIH NeuroBioBank at the University of Maryland, and prepared according to a standardized protocol (http://medschool.umaryland.edu/btbank/method2.asp) under the supervision of the NIH NeuroBioBank ethical guidelines. UMB1465 and UMB4638 died of injuries sustained in motor vehicle accidents, while UMB4643 died of cardiovascular disease. Bulk DNA samples and single neuronal nuclei amplified by multiple displacement amplification (MDA) were prepared and whole-genome sequenced by Illumina HiSeq platforms as part of previous studies in our lab (16). The average sequencing depth was about 40X for single neurons and about 200X for bulk brain samples.

### Estimation of cell-specific dropout rate and error rate

Germline heterozygous mutations were called by GATK HaplotypeCaller (51) from the wholegenome sequencing data from bulk brain DNA samples, and only common SNPs annotated in the 1000 Genome Project (52) were considered to reduce false positive calls. To estimate cellspecific allele dropout rate, we calculate the proportion of germline heterozygous sites that were genotyped as reference-or alternative-homozygous in single-cell sequencing data. One neuron of UMB4643 with significantly lower allele dropout rate (Z-score < −2) was excluded from subsequent analyses, since it likely represented a doublet from FANS sorting. Similarly, we also extracted the reference-homozygous sites at the 3’ adjacent position of each germline heterozygous mutation and calculate the proportion of heterozygous and alternative-homozygous genotypes to estimate the genome-wide error rate in each single-cell.

### Framework of single-cell MosaicHunter

The overall framework of single-cell MosaicHunter (scMH) was illustrated in Fig. 2A. sSNV candidates were first called from the bulk sequencing data using a Bayesian graphical model (20, 21), in which the likelihoods of somatic mutation and three genotypes of inherited mutation were calculated with the consideration of binomial sampling variation and base-calling errors (Fig. 2A, left panel). The presence or absence of somatic mutation in each single-cell was then inferred by adapting the likelihood and allele fraction (*f*) of somatic mutation estimated from bulk sample as prior probability, after controlling the cell-specific allele dropout rate (*d*) and error rate (*e*) (Fig. 2A, right panel). Specifically, the transition matrix between bulk and single-cell genotypes was developed as below,

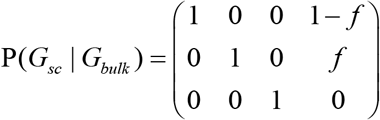

where each column denotes reference-homozygous, heterozygous, alternative-homozygous, and mosaic genotype for bulk sequencing, and each row denotes reference-homozygous, heterozygous, and alternative-homozygous genotype for single-cell sequencing. The genotype likelihoods in single-cells were further adjusted for allele dropout rate (*d*) and error rate (*e*) as below,

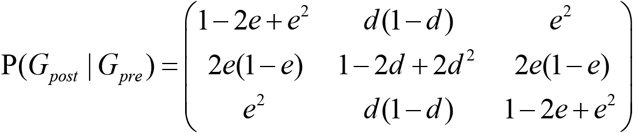

where each column and row denotes reference-homozygous, heterozygous, and alternative-homozygous genotype before and after adjustment for single-cell sequencing. Single-cell genotypes were binarized as mutant or wildtype by comparing the posterior probability of heterozygous genotype to an empirical threshold. For each candidate site, the proportion of mutant cells was calculated to further filter out germline mutations. Candidate sites with >50% cells showing aberrant single-cell allele fractions were also removed to exclude hotspots of technical artifacts. In “bulk-free” mode with synthetic bulk generated from in silico merging sequencing data from multiple single-cells, scMH would only consider sSNVs which were shared by at least two single-cells.

### Somatic SNV calling and performance comparison

Paired-end reads from bulk and single-cell whole-genome sequencing data were aligned to the GRCh37 human reference genome by BWA (53), and then processed by GATK (51) and Picard (http://broadinstitute.github.io/picard/) for the removal of duplicated and error-prone reads, indel realignments, and base-quality recalibrations. sSNVs in neurons of UMB1465, UMB4638, and UMB4643 were called by scMH and four other tools including Monovar (22), SCcaller (23), LiRA (24), and Conbase (25). Sensitivity was estimated as the detected proportion of lineage-informative sSNVs that had been previously identified and validated in these three brain samples (16). False discovery rate (FDR) was measured as the proportion of lineage-informative mutations that were shared by cells from conflicting clades (16).

### Validation of somatic SNVs

Validation of somatic SNVs called from scMH was performed using PCR of 200-500 bp amplicons including the mutated base, followed by Sanger sequencing. All variants were validated in independently sorted single neuronal nuclei amplified by MDA.

### Generation of simulated single-cell whole-genome sequencing data

To estimate the sensitivity of scMH to detect lineage-informative sSNVs, bulk and single-cell sequencing data with varied somatic mutation rates was generated *in silico (SI Appendix,* Fig. S4A). First, we developed a simplistic model to mimic the process of early embryogenesis: 1) ten rounds of symmetric cell division was applied to generate 1024 (2^10^) daughter cells derived from a single zygote, in which somatic mutations was randomly introduced at a rate of 1, 2, 5, or 10 mutations per round; 2) each daughter cells accumulated cell-specific somatic mutations for another ten rounds with the same mutation rate. Then, for each daughter cell, sequencing reads of chromosome 1 was generated at 40X by ART (54) with default parameters for Illumina platforms, and then germline mutations identified from NA12878 and somatic mutations generated by our model was introduced to the sequencing read using BAMSurgeon (55), with an allele dropout rate of 1×10^-2^ per base and MDA amplification rate of 1×10^-7^ per base that were estimated from real single-cell sequencing data. Finally, we randomly selected 80 cells (consistent with the detection threshold of scMH in real brain bulk samples) from the 1024 daughter cells and merged their sequencing data with a down-sampling of 200X to generate the bulk sequencing data, and another 16 cells was randomly selected for benchmarking the performance of scMH. Our simulation data suggested that scMH was able to detect, on average, 67% and 86% of cell-shared sSNVs with PFC bulk or synthetic bulk, respectively *(SI Appendix,* Fig. S4B).

### Design and selection of Taqman genotyping and gene expression probes

Taqman genotyping probes for all validated sSNVs were designed using custom Taqman assay design tool provided Thermo Fisher Scientific. Off-the-shelf Taqman gene expression probes were ordered from Thermo Fisher Scientific. All designed probes were tested by ddPCR using human genomic DNA (Human male, Promega) as a negative control. Gene expression probes were further tested by isolated bulk brain RNA as a positive control. Genotyping probes were also tested by comparing the detected mosaic fractions and the fractions calculated from bulk sequencing (Fig. 3C, D).

### Parallel RNA and DNA analysis after Deep-sequencing (PRDD-seq)

Single nuclei from postmortem brain samples were isolated using fluorescence-activated nuclear sorting (FANS) for NeuN as described previously (56). Isolated single neuronal nuclei were directly sorted into CellsDirect One-Step qRT-PCR (Thermo Fisher Scientific) pre-amplification buffers containing 0.14x Taqman gene expression assays and SNP genotyping assays. Preamplification of all cDNA and genomic DNA amplicons were performed directly after the FANS sorting. Following pre-amplification, samples were diluted 10-fold and loaded onto 96.96 genotyping or 192.24 gene expression dynamic assay integrated fluidic circuits for standard amplification per manufacturer’s instructions (Biomark, Fluidigm). Genotype and gene expression were further determined by Biomark machine and analyzed by Biomark & EP1 software (Fluidigm).

### 10X Genomics preparation and sequencing

Standard 10X Genomics Chromium 3’ (v2 chemistry) was carried out according to the manufacturer’s recommendation. Single nuclei from postmortem brain samples were isolated using FANS for NeuN, and were loaded onto a 10X Genomics Chromium chip. Reverse transcription and library preparation was performed using the 10X Genomics Single Cell v2 kit following the 10X Genomics protocol. The library was then sequenced on one lane of Illumina NextSeq-500 with a high-output kit.

### Single-cell RNA sequencing analysis

The expression matrix of 10X Genomic single-cell RNA sequencing (scRNAseq) was generated by Cell Ranger following the recommended protocols. The expression matrix and cell annotations of SMART-seq-based scRNAseq for human MTG (5) was downloaded from the website (https://celltypes.brain-map.org/rnaseq/). Variance normalization, clustering and visualization were performed by Pagoda2 (57) using a similar protocol to Lake et al (7). Cell clusters containing more than 50 cells were plotted on the t-SNE map, and the annotation of cortical layer (upper, middle, lower) for excitatory neurons and subtypes for inhibitory neurons was manually curated for each cluster according to the expression level of marker genes (*SI Appendix,* Fig. S1 and S2). Considering that Layer 1 dissections of MTG nuclei included the upper part of Layer 2 and the absence of excitatory neurons in the Layer 1 of MTG based on in situ labeling (5), all the MTG Layer 1 excitatory neurons were re-annotated as Layer 2. To further compare the expression profile of cells clusters between two scRNAseq datasets, we calculated the cosine similarity of average expression level for marker genes (*S*) between any two cell clusters. Cell clusters were then hierarchically clustered using the Ward’s method with a distance of 1 – *S*.

### Joint analysis of PRDD-seq and scRNAseq cells

To understand the cell type and cell origin of PRDD-seq cells, we utilized their gene expression profiles to map them onto the t-SNE maps of scRNAseq. PRDD-seq cells were firstly separated into excitatory or inhibitory neurons according to the expression of excitatory or inhibitory marker genes (*SI Appendix,* Table S2), and cells with no or conflicting expression of these marker genes were excluded. For excitatory neurons, missing expression status for layer marker genes (*SI Appendix,* Table S2) were inferred if any layer-specific genes for a given layer were expressed. The cosine similarity matrix was then generated by comparing PRDD-seq cells against scRNAseq cells. For each PRDD-seq cell, its cell cluster was determined by the majority voting among its 25-nearest scRNAseq cells in cosine similarity (Fig. 1I), and the cell type and cortical layer information of PRDD-seq cell was further annotated based on their assigned cell cluster in scRNAseq datasets. To benchmark how accurately we could infer cell type and layer annotation from the 30 marker genes profiled in PRDD-seq cells, we randomly sampled 200 scRNAseq cells from each of the seven cell types (upper, middle, lower layer excitatory neurons and VIP+, SST+, LAMP5+, pPVALB+ inhibitory neurons) from 10X Genomic dataset and each of the six dissected cortical layers from SMART-seq dataset, and only extracted the expression profiles of 30 marker genes from each scRNAseq cell. Using the same majority voting strategy, we assigned them back to cell clusters on the t-SNE map. As shown in Fig. 3D, the majority of the randomly sampled cells can be correctly assigned to their original cell type and layer annotation, suggesting the accuracy of our mapping strategy in PRDD-seq.

### Quantification and statistical analysis

All data are reported as mean ± 95% confident interval (CI) unless mentioned otherwise. All of the statistical details can be found in the figure legends, figures, and Results. Significance was defined for p values smaller than 0.05. All tests were performed using the R software package (version 3.5.0).

### Data and code availability

Sequencing data was deposited in the NCBI SRA with accession numbers SRP041470 and SRP061939. MosaicHunter is publicly available at http://mosaichunter.cbi.pku.edu.cn/. Config files of single-cell MosaicHunter (scMH) and other scripts about PRDD-seq can be accessed at https://github.com/AugustHuang/PRDD-seq.

## Author Contributions

A.Y.H. and P.L. conceived the project and C.A.W. supervised it. A.Y.H. and P.L. developed the scMH algorithm. P.L. developed PRDD-seq and performed experiments. A.Y.H. performed computational and statistical analyses. R.E.R., S.N.K., and Y.D. helped with validation of sSNVs. C.J.K. helped with and provided insight on the comparison of PRDD-seq and scWTA. S.K.A. assisted with interpretation of neurodevelopmental discoveries. R.D.H., T.E.B., J.M. and E.S.L. generated the MTG single-cell RNA sequencing data, and provided it prior to publication. E.A.L. and P.J.P. provided suggestions on computational analyses. A.Y.H. and P.L. wrote the manuscript supervised by C.A.W., with input from all other authors.

## Acknowledgements

We thank R. Mattieu, K. Brownstein, J. Li, Flow Cytometry Facility in Boston Children’s Hospital, BCH IDDRC Molecular Genetics Core Facility, and the Research Computing group at Harvard Medical School for assistance. We thank G. Fishell, C. Harwell, and F. Vaccarino for comments on the manuscript. Human tissue was obtained from the NIH NeuroBioBank at the University of Maryland and Autism BrainNet, and we thank the donors and their families for their invaluable donations for the advancement of science. P.L is a Howard Hughes Medical Institute – Helen Hay Whitney Foundation Fellow. C.A.W. is supported by the Manton Center for Orphan Disease Research, the Allen Discovery Center program through The Paul G. Allen Frontiers Group, grants from the NINDS (R01NS032457 and U01MH106883), and grant U01MH106883 from the NIMH. C.A.W. is an Investigator of the Howard Hughes Medical Institute.

