## Supplementary Information for "Parallel RNA and DNA analysis after Deep-sequencing (PRDD-seq) reveals cell type specific lineage patterns in human brain"

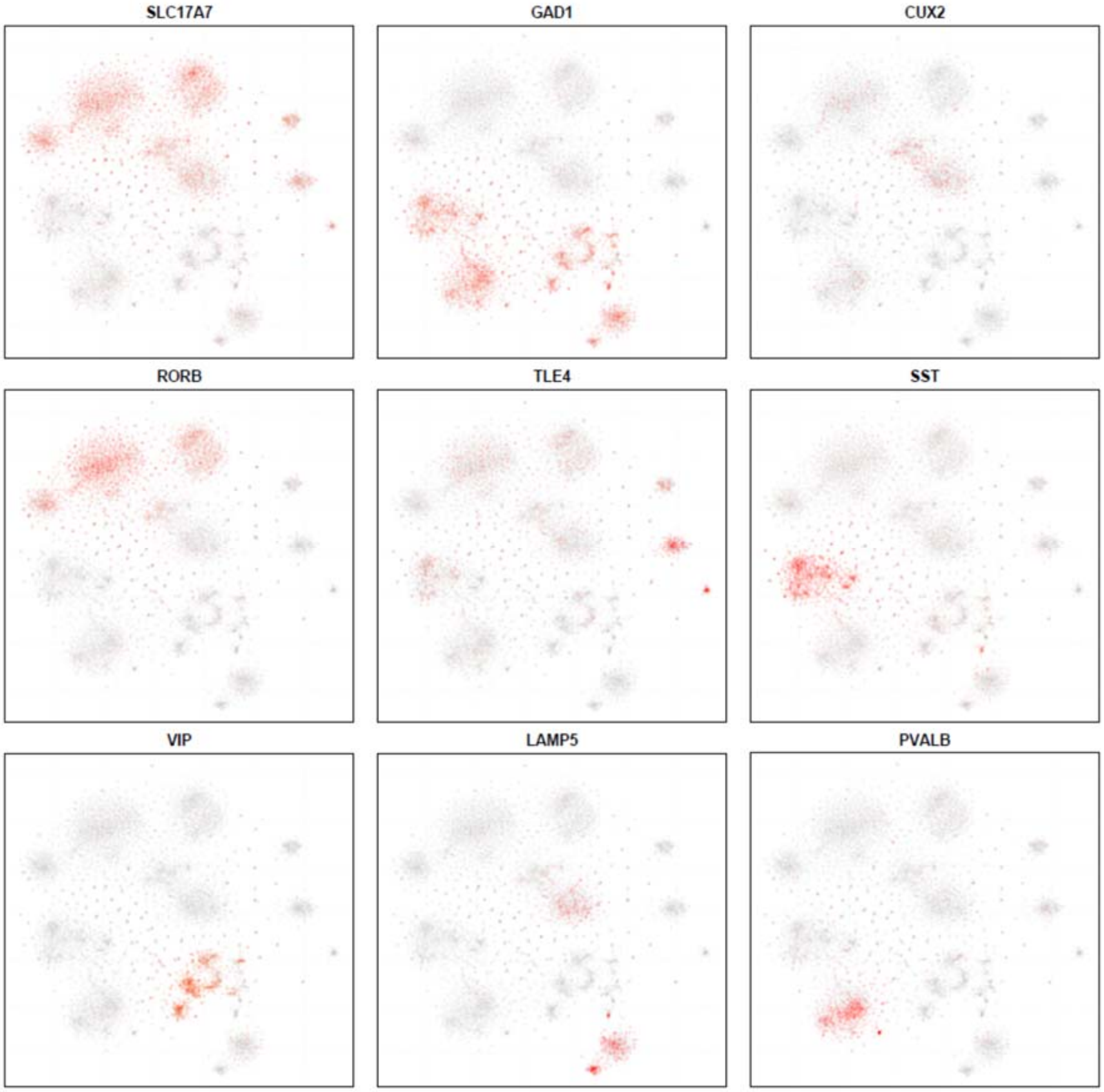


**Figure S1. t-SNE plots for the single-neuron expression levels of marker genes in 10X Genomics scRNAseq dataset.** SLC17A7: excitatory neuron; GAD2: inhibitory neuron; CUX2: upper layer excitatory neuron; RORB: middle layer excitatory neuron; TLE4: lower layer excitatory neuron; SST, VIP, LAMP5, and PVALB: four different subtypes of inhibitory neuron.

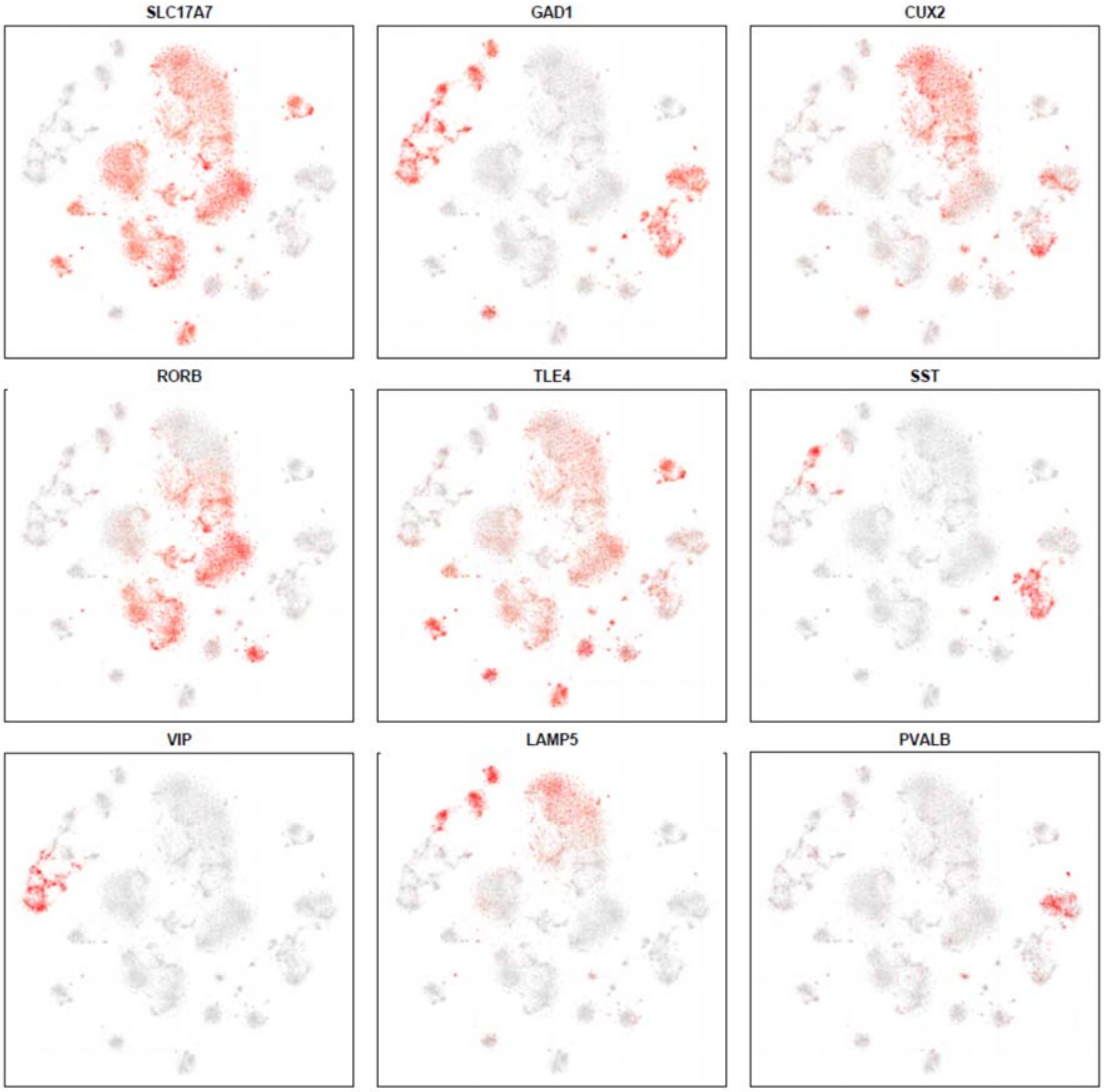


**Figure S2. t-SNE plots for the single-neuron expression levels of marker genes in SMART-seq scRNAseq dataset.** SLC17A7: excitatory neuron; GAD2: inhibitory neuron; CUX2: upper layer excitatory neuron; RORB: middle layer excitatory neuron; TLE4: lower layer excitatory neuron; SST, VIP, LAMP5, and PVALB: four different subtypes of inhibitory neuron.

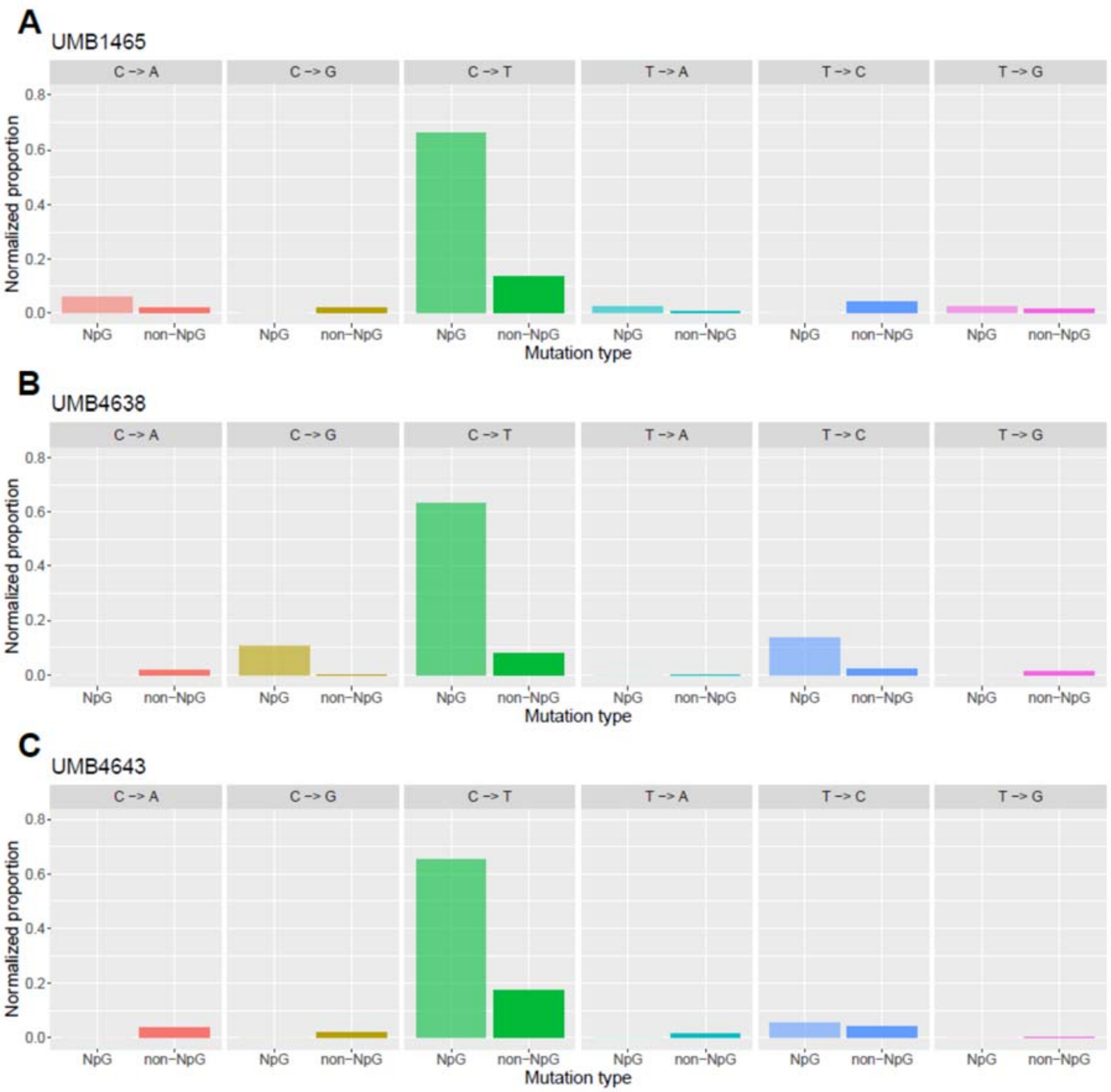


**Figure S3. Mutation spectrum of validated lineage-informative sSNVs in UMB1465 (A), UMB4638 (B), and UMB4643 (C).** The sSNVs showed enrichment for C>T mutations and especially in CpG sites.

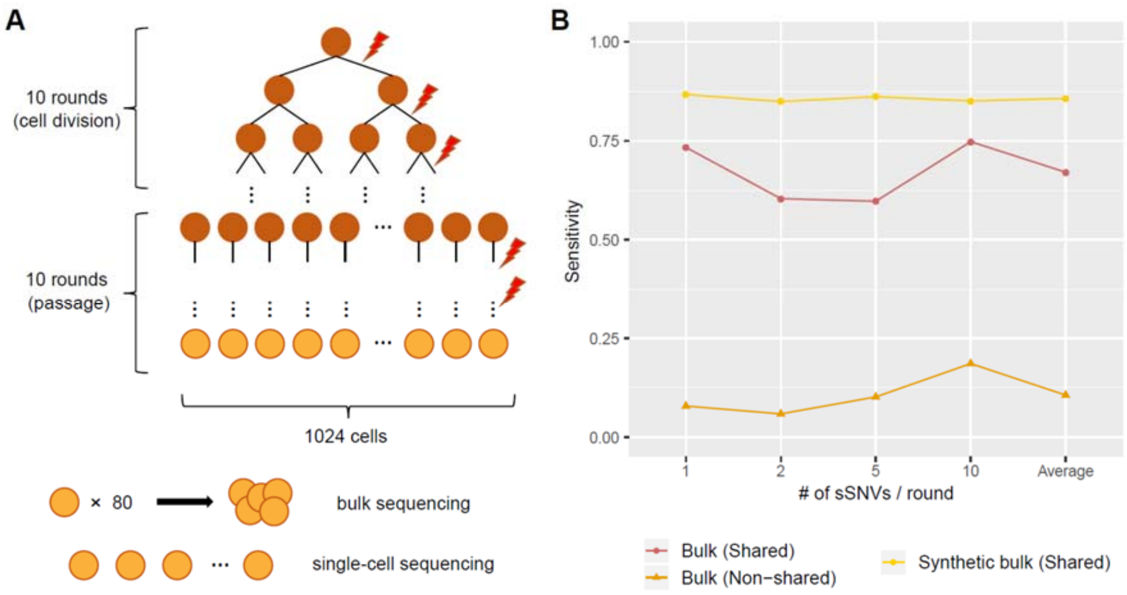


**Figure S4. Performance of scMH benchmarked by simulated sequencing data with varied somatic mutation rates.** A. Generation of simulated single-cell and bulk sequencing data. For each simulation, 1024 single-cells were derived from a single zygote after ten rounds of cell division followed by another ten rounds for the accumulation of cell-specific mutations, with 1, 2, 5, or 10 random somatic mutations introduced per round. B. Sensitivity for detecting shared and non-shared sSNVs by two modes of scMH.

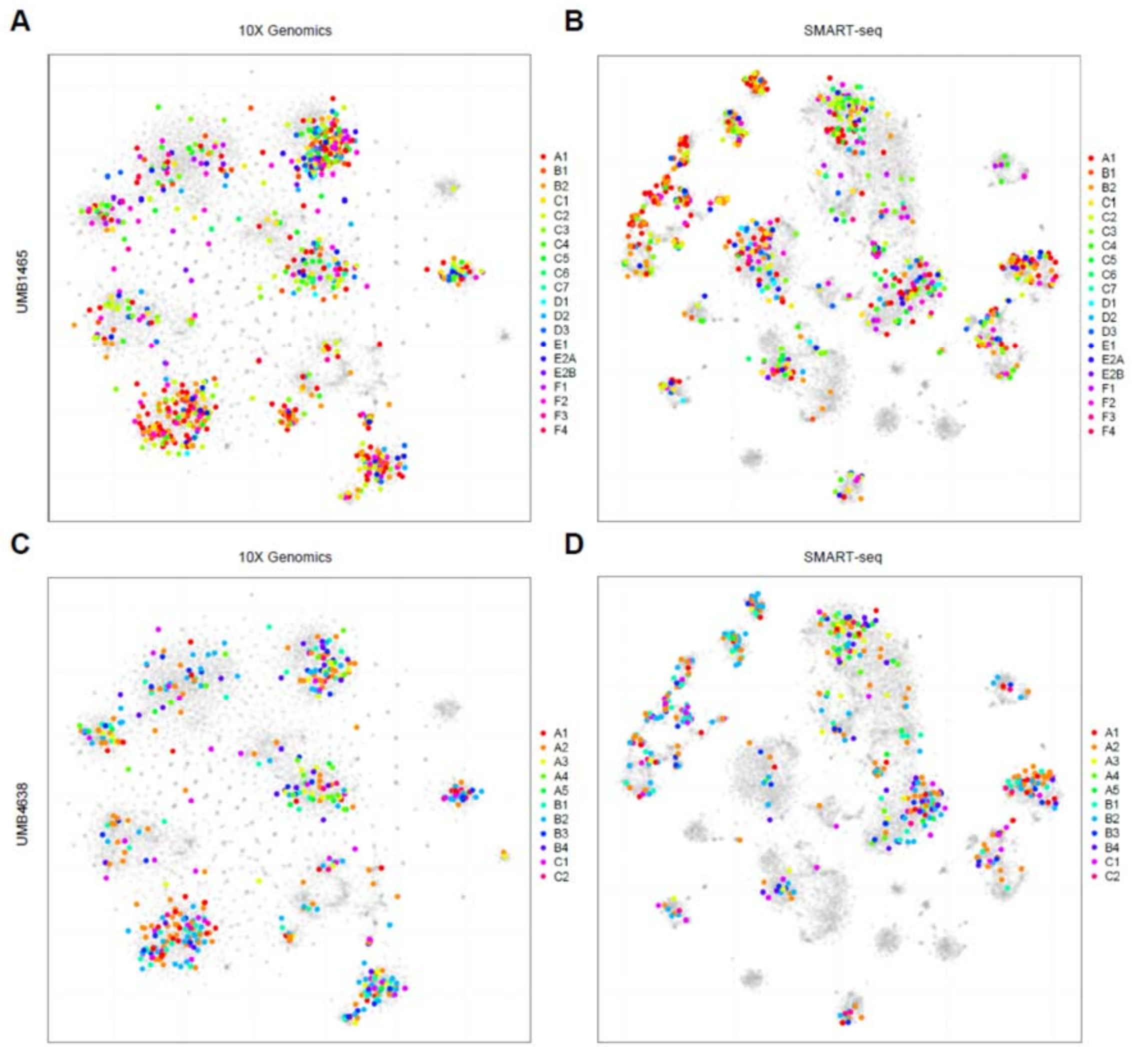


**Figure S5. Cell type classification of PRDD-seq cells onto human brain scRNAseq t-SNE plots.** A-B. PRDD-seq cells from UMB1465 onto 10X Genomics (A) and SMART-seq (B) datasets. C-D. PRDD-seq cells from UMB1465 onto 10X Genomics (C) and SMART-seq (D) datasets. PRDD-seq cells were assigned to different t-SNE clusters based on the expression profile similarity of 30 marker genes, and cells with different lineage markers were labeled by different colors.

**Table S1. Validated sSNVs used in lineage tree reconstruction**

| Individual | ID | Chr | Position | Ref allele | Alt allele | Mosaic fraction in PRDD-seq cells |
|---|---|---|---|---|---|---|
| UMB1465 | A1 | 4 | 171901716 | C | T | 18.07% |
| UMB1465 | B1-1 | 9 | 76779133 | T | G | 17.94% |
| UMB1465 | B1-2 | 11 | 73504764 | G | C | 17.94% |
| UMB1465 | B2-1 | 7 | 25690508 | G | A | 12.05% |
| UMB1465 | B2-2 | 11 | 40316580 | C | T | 12.05% |
| UMB1465 | C1-1 | 8 | 135332005 | A | C | 33.20% |
| UMB1465 | C1-2 | 5 | 3926977 | C | G | 33.20% |
| UMB1465 | C2-1 | 5 | 156493734 | T | C | 28.11% |
| UMB1465 | C2-2 | 3 | 20458612 | C | T | 28.11% |
| UMB1465 | C3 | 3 | 18578000 | G | T | 17.27% |
| UMB1465 | C4-1 | 7 | 17623547 | C | T | 9.10% |
| UMB1465 | C4-2 | 6 | 4936481 | G | A | 9.10% |
| UMB1465 | C5-1 | 2 | 84862977 | C | T | 4.42% |
| UMB1465 | C5-2 | 14 | 76355049 | C | A | 4.42% |
| UMB1465 | C6 | 7 | 93952235 | A | T | 2.28% |
| UMB1465 | C7 | X | 138790531 | G | A | 0.67% |
| UMB1465 | D1 | 4 | 170816788 | G | A | 9.24% |
| UMB1465 | D2 | 16 | 62830915 | C | T | 7.50% |
| UMB1465 | D3 | 7 | 15742416 | G | A | 4.42% |
| UMB1465 | E1 | 9 | 4449186 | G | A | 5.49% |
| UMB1465 | E2A-1 | 8 | 40724674 | G | A | 2.68% |
| UMB1465 | E2A-2 | 13 | 109550610 | G | A | 2.68% |
| UMB1465 | E2B | 9 | 8100716 | T | C | 1.47% |
| UMB1465 | F1-1 | X | 135442114 | T | G | 14.59% |
| UMB1465 | F1-2 | 4 | 154649455 | C | T | 14.59% |
| UMB1465 | F1-3 | 8 | 50806767 | T | A | 14.59% |
| UMB1465 | F2-1 | 18 | 4149577 | G | T | 11.24% |
| UMB1465 | F2-2 | 7 | 68820962 | G | A | 11.24% |
| UMB1465 | F3 | 12 | 52644508 | C | T | 1.61% |
| UMB1465 | F4 | 2 | 26914023 | G | A | 0.40% |
| UMB4638 | A1 | 3 | 55954310 | G | T | 46.04% |
| UMB4638 | A2-1 | 6 | 47874596 | C | T | 38.96% |
| UMB4638 | A2-2 | 6 | 127381695 | A | C | 38.96% |
| UMB4638 | A3 | 2 | 167813464 | G | A | 12.29% |
| UMB4638 | A4 | 22 | 27399001 | A | G | 6.04% |
| UMB4638 | A5-1 | 17 | 48665916 | G | A | 2.29% |
| UMB4638 | A5-2 | 15 | 22394229 | C | T | 2.29% |
| UMB4638 | B1 | 8 | 14156088 | C | T | 42.29% |

| | | | | | | |
|---|---|---|---|---|---|---|
| UMB4638 | B2 | 3 | 25001431 | C | G | 31.25% |
| UMB4638 | B3 | 11 | 106806293 | C | T | 11.67% |
| UMB4638 | B4 | 5 | 173266954 | G | A | 5.42% |
| UMB4638 | C1 | 19 | 47565457 | C | T | 11.67% |
| UMB4638 | C2 | 3 | 11688960 | G | A | 3.13% |

**Table S2. Information of biomarkers used in this study**

| Gene | Taqman Probe ID | Entrez ID | Gene description | Note |
|---|---|---|---|---|
| BHLHE22 | hs010844964_s1 | 27319 | basic helix-loop-helix family member e22 | middle layer |
| CCK | hs00174937_m1 | 885 | cholecystokinin | inhibitory |
| CUX2 | hs00322624_m1 | 23316 | cut like homeobox 2 | upper layer |
| CXCL14 | hs01557413_m1 | 9547 | C-X-C motif chemokine ligand 14 | |
| FOXP2 | hs00362818_m1 | 93986 | forkhead box P2 | neuronal |
| GAD1 | hs01065893_m1 | 2571 | glutamate decarboxylase 1 | inhibitory |
| GAD2 | hs00609534_m1 | 2572 | glutamate decarboxylase 2 | inhibitory |
| GFAP | hs00909233_m1 | 2670 | Glial Fibrillary Acidic Protein | |
| GAPDH | hs02786624_g1 | 2597 | glyceraldehyde-3-phosphate dehydrogenase | housekeeping |
| GLRA3 | hs00923566_m1 | 8001 | glycine receptor alpha 3 | upper layer |
| KCNK2 | hs01005159_m1 | 3776 | potassium two pore domain channel subfamily K member 2 | |
| LAMP5 | hs00202136_m1 | 24141 | lysosomal associated membrane protein family member 5 | upper layer; inhibitory - LAMP5 |
| NPY | hs00173470_m1 | 4852 | neuropeptide Y | |
| OLIG1 | hs00744293_s1 | 116448 | Oligodendrocyte Transcription Factor 1 | |
| NR4A2 | hs01117527_g1 | 4929 | nuclear receptor subfamily 4 group A member 2 | lower layer |
| PAX6 | hs01088114_m1 | 5080 | paired box 6 | |
| RBFOX3 | hs01370654_m1 | 146713 | RNA binding fox-1 homolog 3 | neuronal |
| RBP4 | hs00924047_m1 | 5950 | retinol binding protein 4 | middle layer |
| RELN | hs01022646_m1 | 5649 | reelin | |
| RORB | hs00199445_m1 | 6096 | RAR related orphan receptor B | middle layer |
| SLC17A6 | hs00220439_m1 | 57084 | solute carrier family 17 member 6 | excitatory |
| SLC17A7 | hs00220404_m1 | 57030 | solute carrier family 17 member 7 | excitatory |
| SLC32A1 | hs00369773_m1 | 140679 | solute carrier family 32 member 1 | inhibitory |
| SOX2 | hs01053049_s1 | 6657 | SRY-box 2 | |
| SOX6 | hs00264525_m1 | 55553 | SRY-box 6 | |
| SST | hs00356144_m1 | 6750 | somatostatin | inhibitory - SST |
| SYNPR | hs01548398_m1 | 132204 | synaptoporin | neuronal |
| TBR1 | hs00232429_m1 | 10716 | T-box, brain 1 | |
| TLE4 | hs00925588_g1 | 7091 | TLE family member 4, transcriptional corepressor | lower layer |
| VIP | hs00175021_m1 | 7432 | vasoactive intestinal peptide | inhibitory - VIP |

**Table S3. Comparison of excitatory-inhibitory neuron ratio in this and other studies**

| Study | Brain region | Sorting | NeuN staining | Prep | Ex% | Note |
|---|---|---|---|---|---|---|
| Darmanis et al, 2015 | Anterior temporal lobe | No | No | Fluidigm | 62% | fresh whole cells |
| Lake et al, 2016 | PFC and other regions | Yes | Yes | Fluidigm | 70% | |
| Habib et al, 2017 | PFC and other regions | No | No | Drop-seq | 70% | |
| Lake et al, 2018 | PFC and other regions | No | No | Drop-seq | 68% | |
| Hodge et al, 2019 | MTG | Yes | Yes | SMART-seq | 71% | |
| This study | PFC | Yes | Yes | 10X Genomics | 56% | UMB1465 |
| This study | PFC | Yes | Yes | PRDD-seq | 54% | UMB1465&4638 |
